# Severe deficiency of voltage-gated sodium channel Na_V_1.2 elevates neuronal excitability in adult mice

**DOI:** 10.1101/2021.02.02.429384

**Authors:** Jingliang Zhang, Xiaoling Chen, Muriel Eaton, Shirong Lai, Anthony Park, Talha S. Ahmad, Jiaxiang Wu, Zhixiong Ma, Zhefu Que, Ji Hea Lee, Tiange Xiao, Yuansong Li, Yujia Wang, Maria I. Olivero-Acosta, James A. Schaber, Krishna Jayant, Zhuo Huang, Nadia A. Lanman, William C. Skarnes, Yang Yang

## Abstract

*Scn2a* encodes voltage-gated sodium channel Na_V_1.2, which mediates neuronal firing. The current paradigm suggests that Na_V_1.2 gain-of-function variants enhance neuronal excitability resulting in epilepsy, whereas Na_V_1.2 deficiency impairs neuronal excitability contributing to autism. In this paradigm, however, why about a third of patients with Na_V_1.2 deficiency still develop seizures remains a mystery. Here we challenge the conventional wisdom, reporting that neuronal excitability is increased with severe Na_V_1.2 deficiency. Using a unique gene-trap knockout mouse model of *Scn2a*, we found enhanced intrinsic excitabilities of principal neurons in the cortico-striatal circuit, known to be involved in *Scn2a*-related seizures. This increased excitability is autonomous, and is reversible by genetic restoration of *Scn2a* expression in adult mice. Mechanistic investigation reveals a compensatory downregulation of potassium channels including K_V_1.1, which could be targeted to alleviate neuronal hyperexcitability. Our unexpected findings may explain Na_V_1.2 deficiency-related epileptic seizures in humans and provide molecular targets for potential interventions.

**TEASER:** Severe Na_V_1.2 deficiency results in neuronal hyperexcitability via the compensatory downregulation of potassium channels.

**HIGHLIGHTS:**

1. Severe Na_V_1.2 deficiency results in enhanced excitability of medium spiny neurons (MSNs) and pyramidal neurons in adult mice;
2. Increased neuronal excitability in MSNs is accompanied by elevated voltage threshold;
3. Na_V_1.2 deficiency-related hyperexcitability is reversible with the restoration of *Scn2a* expression, and is autonomous;
4. The expression of the K_V_1.1 channel has a compensatory reduction in neurons with Na_V_1.2 deficiency, and KV channels openers normalize the neuronal excitability;
5. The enhanced excitability in brain slices translates to elevated *in vivo* firing commonly associated with seizures.

## INTRODUCTION

NaV1.2 channel, encoded by *SCN2A*, is a major voltage-gated sodium channel expressed in the central nervous system (CNS) supporting the action potentials (AP) firing (*1*) (*2*). NaV1.2 is strongly expressed in the principal neurons of the cortico-striatal circuit, including pyramidal neurons of the medial prefrontal cortex (mPFC) and medium spiny neurons (MSNs) of the caudate-putamen (CPu) in the striatum (*3–5*). Gain-of-function (GoF) variants of *SCN2A* are closely associated with epileptic seizures, whereas loss-of-function (LoF) or protein-truncating variants of *SCN2A* (collectively referred to as NaV1.2 deficiency) are leading genetic causes of autism spectrum disorder (ASD) and intellectual disability (ID) (*6–10*). The conventional paradigm suggests that GoF variants of *SCN2A* increase the excitability of principal neurons resulting in epilepsy, whereas NaV1.2 deficiency impairs the excitability of principal neurons leading to ASD (*2*). However, clinical studies found that a significant portion of patients with NaV1.2 deficiency develop “late-onset” intractable seizures (*11, 12*). As hyperexcitability and hypersynchronization of neuronal firings are suggested as the basis of seizures (*13*), it is thus intriguing how NaV1.2 deficiency, predicted to reduce neuronal excitability, contributes to epileptic seizures.

To understand NaV1.2 deficiency-related pathophysiology, mouse models were generated. Homozygous *Scn2a*^*−/−*^ knockout mice die perinatally (*14, 15*); Heterozygous *Scn2a*^*+/−*^ mice (with ~50% NaV1.2 expression level) survive to adulthood, but the earlier study did not find notable abnormalities in *Scn2a*^*+/−*^ mice (*14*). More recently, absence-like seizures were reported in adult male *Scn2a*^*+/−*^ mice (*16*). It is suggested that the CPu of the striatum and the mPFC of the cortex are key brain regions in which absence seizure-like spike-wave discharges (SWDs) were identified (*16, 17*). Indeed, the cortico-striatal circuit is highly involved in ASD as well as seizures, and the excitability of principal neurons in this circuit could strongly influence seizure susceptibility (*18, 19*). Despite these *in vivo* findings, recordings in brain slices, however, revealed unchanged AP firings and reduced excitatory postsynaptic current in pyramidal neurons of adult *Scn2a*^*+/−*^ mice (*16, 20*), leaving cellular mechanisms ambiguous.

It is not uncommon that phenotypes observed in hemizygous patients do not manifest in heterozygous mouse models. In fact, it is known that mice are more tolerant than humans to certain gene expression reduction (*21*). Therefore, the heterozygous knockout with a close to 50% reduction in *Scn2a* protein level may not be sufficient to render major phenotypes in mice (*21*). A more substantial reduction of gene expression could be essential to produce robust phenotypes in the mouse model of NaV1.2 deficiency. Because *Scn2a* null (100% knockout) is lethal, we thus generated a novel NaV1.2-deficient mouse model via a gene-trap knockout (gtKO) strategy (*22*). These mice display many behavioral abnormalities, modeling aspects of phenotypes in humans with *SCN2A* deficiency (*22*). Using this unique mouse model, we investigated how severe NaV1.2 deficiency affects neuronal excitabilities of principal neurons in the cortico-striatal circuit. Our results demonstrate a surprising hyperexcitability phenotype in neurons, in which the compensatory downregulation of the potassium channels is likely to be an underlying mechanism.

## RESULTS

### Neurons expressing substantially low NaV1.2 exhibit elevated excitability

To understand how a severe NaV1.2 deficiency affects the function of neurons, we utilized a gene-trap knockout mouse model of *Scn2a*. Homozygous *Scn2a*^*gtKO/gtKO*^ mice (referred to as HOM herein) can survive to adulthood, and have a substantial reduction of NaV1.2 expression (~25% of the WT level) (*22*). Because the gene-trap cassette contains a *LacZ* element, which is driven by the native NaV1.2 promoter (**Figure S1A**) (*23, 24*), we used *LacZ*-staining as a surrogate to determine the expression and distribution of NaV1.2 in the brain. Our data showed that *Scn2a* is widely expressed in the mouse brain including the cortex and striatum (**Figure S1B**), which is consistent with previous studies of *Scn2a* distribution (*3–5*).

The CPu is a common node for ASD and seizures, and is one of the major brain regions involved in the *Scn2a*-related absence-like seizures (*16–18*). Previous study and our *LacZ*-staining suggested that NaV1.2 is highly expressed in the CPu. To further confirm these results, we performed Western blot analysis. We found that the heterozygous (HET) *Scn2a*^*WT/gtKO*^ mice have ~60% of WT NaV1.2 protein level in the CPu tissues, whereas the homozygous (HOM) *Scn2a*^*gtKO/gtKO*^ mice have a much lower level at 34% (**Figure S1C**). This result is largely consistent with our initial characterization of this mouse model using whole-brain samples (*22*). To understand how a severe deficiency of NaV1.2 affects neuronal excitability, we performed *ex vivo* patch-clamp recordings in brain slices from adult *Scn2a*^*gtKO/gtKO*^ mice. Unexpectedly, we found that the striatal principal medium spiny neurons (MSNs) from *Scn2a*^*gtKO/gtKO*^ mice were markedly more excitable (**Figure 1A-C**). The current-injection triggered action potential (AP) number was significantly elevated in MSNs from *Scn2a*^*gtKO/gtKO*^ mice compared to WT littermates. We also observed depolarized resting membrane potential (RMP) and increased input resistance of these MSNs (**Figure 1D, E**), which were in line with the increased neuronal excitability. Phase-plane plot analysis showed that the AP waveform in *Scn2a*^*gtKO/gtKO*^ mice was altered (**Figure 1F, G**). While rheobase was reduced, interestingly we detected a higher voltage threshold, reduced AP amplitude, elevated fast after-hyperpolarization (fAHP), and increased half-width values in MSNs from *Scn2a*^*gtKO/gtKO*^ mice (**Figure 1H-L**). Voltage-dependent conductance can affect neuronal RMP (*25*), and RMP is known to influence neuronal excitability (*26*). We thus performed recordings at a fixed membrane potential (MP) to understand whether the altered RMP is a major factor for this observed hyperexcitability of MSNs. However, even at the fixed MP, we were still able to detect the enhanced excitability along with the altered AP waveforms in *Scn2a*^*gtKO/gtKO*^ mice (**Figure S1D-M**), suggesting that besides the RMP, other factors are playing essential roles contributing to the neuronal hyperexcitability. Taken together, our data suggest a counterintuitive finding that severe deficiency of NaV1.2 renders an increased (rather than conventionally suggested as decreased) neuronal excitability.

**Fig. 1.**
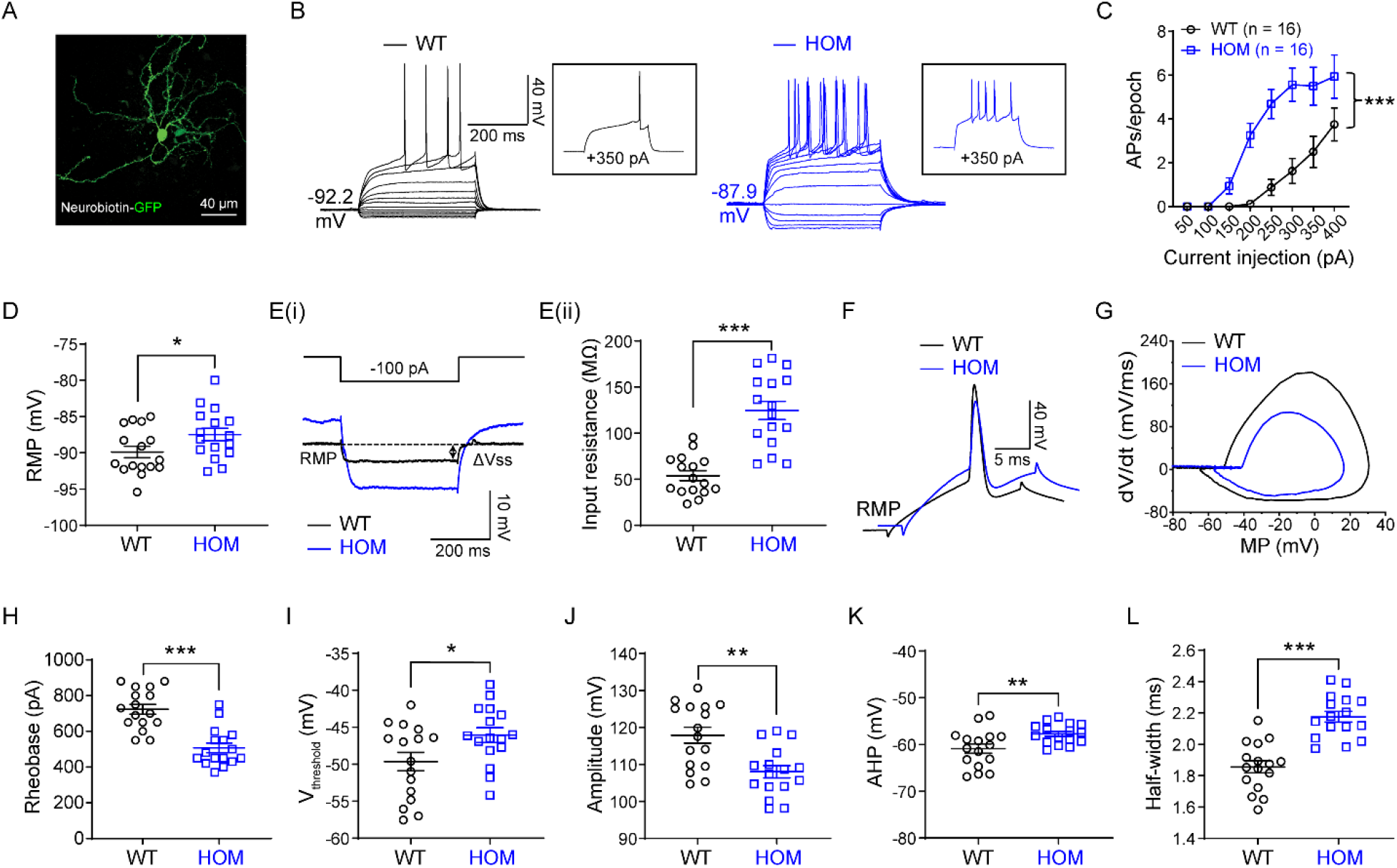
Elevated neuronal firings of striatal medium spiny neurons (MSNs) in adult Na_V_1.2-deficient mice. (**A**) A typical MSN labeled by neurobiotin. Scale bar, 40 μm. (**B**) Representative current-clamp recordings of MSNs from WT (black) and homozygous (HOM), *Scn2a*^*gtKO/gtKO*^ (blue) mice were obtained at the resting membrane potential (RMP). A series of 400-ms hyperpolarizing and depolarizing steps in 50-pA increments were applied to produce the traces. Inset: representative trace in response to 350 pA positive current injection. (**C**) The average number of action potentials (APs) generated in response to depolarizing current pulses. Unpaired two-tailed non-parametric Mann-Whitney *U*-test for each current pulse: ***p < 0.001. (**D**) Individuals and mean RMP values. Unpaired two-tailed Student’s *t*-test: *p < 0.05. (**Ei**) Representative traces in response to 100 pA negative current injection. V_steady-state_ (V_ss_) is the voltage recorded at 0-10 ms before the end of the stimulus. (**Eii**) Individuals and mean input resistance values at the RMP. Unpaired two-tailed Student’s *t*-test: ***p < 0.001. (**F**) Typical spikes of MSNs from WT (black) and HOM (blue) mice were obtained at the normal RMP. (**G**) Associated phase-plane plots. (**H-L**) Individuals and mean spike rheobase, voltage threshold, amplitude, fAHP (fast after-hyperpolarization), and half-width values. Unpaired two-tailed Student’s *t*-test: *p < 0.05; **p < 0.01; ***p < 0.001. Data were shown as mean ± SEM.

### Enhanced excitability is reversible in adult Na_v_1.2-deficient mice with the restoration of *Scn2a* expression, and is autonomous

*Scn2a*^*gtKO/gtKO*^ mice, generated via a gene-trap strategy, has a built-in genetic “rescue” element for manipulations (*24, 27*). The inserted “tm1a” trapping cassette is flanked with *Frt* sites, which can be removed via a flippase recombinase (Flp) to achieve a “tm1c” allele in a temporally and spatially controlled manner (*24*) (**Figure S1A**). This “tm1c” allele is practically a “rescue” allele to restore the expression of the target gene. We performed experiments to restore the *Scn2a* expression by adeno-associated virus (AAV) delivery of codon-optimized Flp (FlpO), with a goal to determine the reversibility of these enhanced neuronal firings in adult mice. Using a PHP.eB.AAV vector, which can be administered via systemic delivery (**Figure 2A**) to transduce neurons across the brain (*28*), we studied the *LacZ* signals (**Figure 2B**) and the protein expression level of *Scn2a*. We found that the FlpO treatment resulted in a partial but significant elevation of Na_v_1.2 protein expression in adult *Scn2a*^*gtKO/gtKO*^ mice compared to control PHP.eB.AAV transduction (**Figure 2C**). Remarkably, this partial restoration of *Scn2a* expression in adult mice translated into changes in neuronal excitability. We found that the adult *Scn2a*^*gtKO/gtKO*^ mice transduced with the AAV-FlpO displayed decreased neuronal excitability of striatal MSNs (**Figure 2D-E)**. In the FlpO-treated group, the triggered AP firing of MSNs in *Scn2a*^*gtKO/gtKO*^ mice was reduced to the WT range, together with the correction of other parameters including the RMP, AP waveform among others (**Figure 2D-J)**. Collectively, our data show that even with a partial restoration of *Scn2a* expression to ~50-60% of WT expression level, we are still able to achieve an almost full rescue of neuronal excitability in adult mice.

**Fig. 2.**
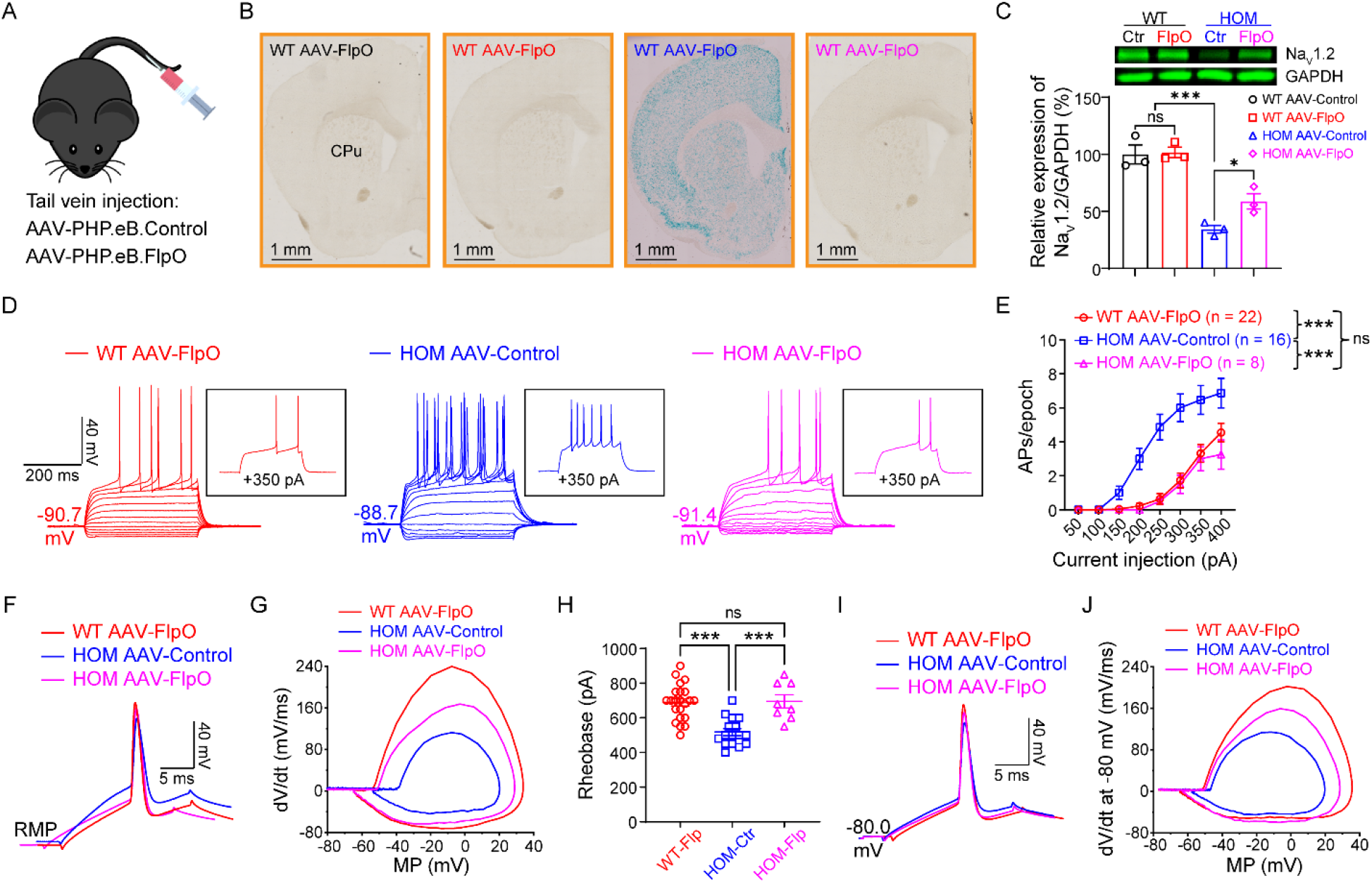
Elevated neuronal firing is reversible by FlpO-mediated restoration of Na_v_1.2 expression in adult Na_v_1.2-deficient mice. (**A**) Cartoon illustration of mice systemically administrated with PHP.eB.AAV-control or AAV-FlpO via tail vein injection. (**B**) Coronal views of *LacZ* staining of striatum from WT and *Scn2a*^*gtKO/gtKO*^ (HOM) mice injected with AAV-control or AAV-FlpO. Blue staining of HOM mice largely disappeared in the AAV-FlpO group. CPu, caudate nucleus and the putamen (dorsal striatum). (**C**) The Western blot analysis showed Na_v_1.2 protein levels in whole-brain tissues from *Scn2a*^*gtKO/*gtKO^ (HOM) mice in AAV-Control or AAV-FlpO group. One-way ANOVA with Bonferroni’s multiple-comparison test: ns, no significance, p > 0.05; *p < 0.05; ***p < 0.001. (**D**) Representative current-clamp recordings of MSNs from WT mice transduced with AAV-FlpO (red), HOM mice transduced with AAV-Control (blue), and HOM mice transduced with AAV-Control (magenta) obtained at the RMP. A series of 400-ms hyperpolarizing and depolarizing steps in 50-pA increments were applied to produce the traces. Inset: representative trace in response to 350 pA positive current injection. (**E**) The average number of APs generated in response to depolarizing current pulses at the RMP. Unpaired two-tailed non-parametric Mann-Whitney *U*-test for each current pulse: ns, no significance, *p > 0.05; ***p < 0.001. (**F**) Typical spikes of MSNs from WT transduced with AAV-FlpO (red), HOM transduced with AAV-Control (blue) and HOM transduced with AAV-Control (magenta) were obtained at the normal RMP. (**G**) Associated phase-plane plots. (**H**) Individuals and average spike rheobase. Unpaired two-tailed Student’s *t*-test: ns, no significance, p > 0.05; ***p < 0.001. (**I**) Typical spikes of MSNs from WT mice transduced with AAV-FlpO (red), HOM mice transduced with AAV-Control (blue), and HOM mice transduced with AAV-Control (magenta) at a fixed membrane potential of −80 mV. (**J**) Associated phase-plane plots at −80 mV. Data were shown as mean ± SEM.

In the cortico-striatal circuit, principal pyramidal neurons of the mPFC project to the striatum, and are suggested to be involved in seizure initiation. As the mPFC is also implicated in the absence-like seizures of *Scn2a*^*+/−*^ mice (*16, 17*), we studied the excitability of layer V pyramidal neurons of the mPFC. We found that the excitability of these neurons was increased significantly compared to the WT mice, and can be reversed by FlpO mediated partial restoration of *Scn2a* expression as well (**Figure S2**). Together, our data suggest that the Na_v_1.2 deficiency-related hyperexcitability exists along the cortico-striatal circuit, manifested in the principal neurons of both cortex and striatum brain regions.

The hyperexcitability seen in neurons with Na_v_1.2 deficiency could come from the altered intrinsic properties independent of other neurons (autonomous), or a result of a disrupted circuit. To distinguish these possibilities, we performed AAV injections of FlpO-mCherry to transduce only a few MSNs in the CPu sparsely. We then performed patch-clamp recordings on adjacent neurons with or without fluorescence (AAV-negative/non-transduced neurons versus AAV-positive/transduced neurons) **(Figure 3A**). Strikingly, our data showed that the transduced neurons (showing fluorescence) display greatly decreased neuronal excitability, compared to non-transduced neurons (showing non-fluorescence) in the same brain slices. In particular, we found that the RMP, input resistance, and the altered AP waveform were reversed in FlpO-transduced neurons of *Scn2a*^*gtKO/gtKO*^ mice (**Figure 3B-L**). Moreover, when we performed the recordings at a fixed membrane potential of −80 mV, similar findings could still be obtained (**Figure S3A-J**). On the other hand, non-transduced neurons displayed hyperexcitability similarly to neurons from *Scn2a*^*gtKO/gtKO*^ mice without virus transduction. Our data indicate that the hyperexcitability of each MSN can be modulated by the expression level of *Scn2a* autonomously, and the Na_v_1.2 deficiency-related hyperexcitability is the intrinsic property of a particular neuron independent of its surrounding neurons or circuit.

**Fig. 3.**
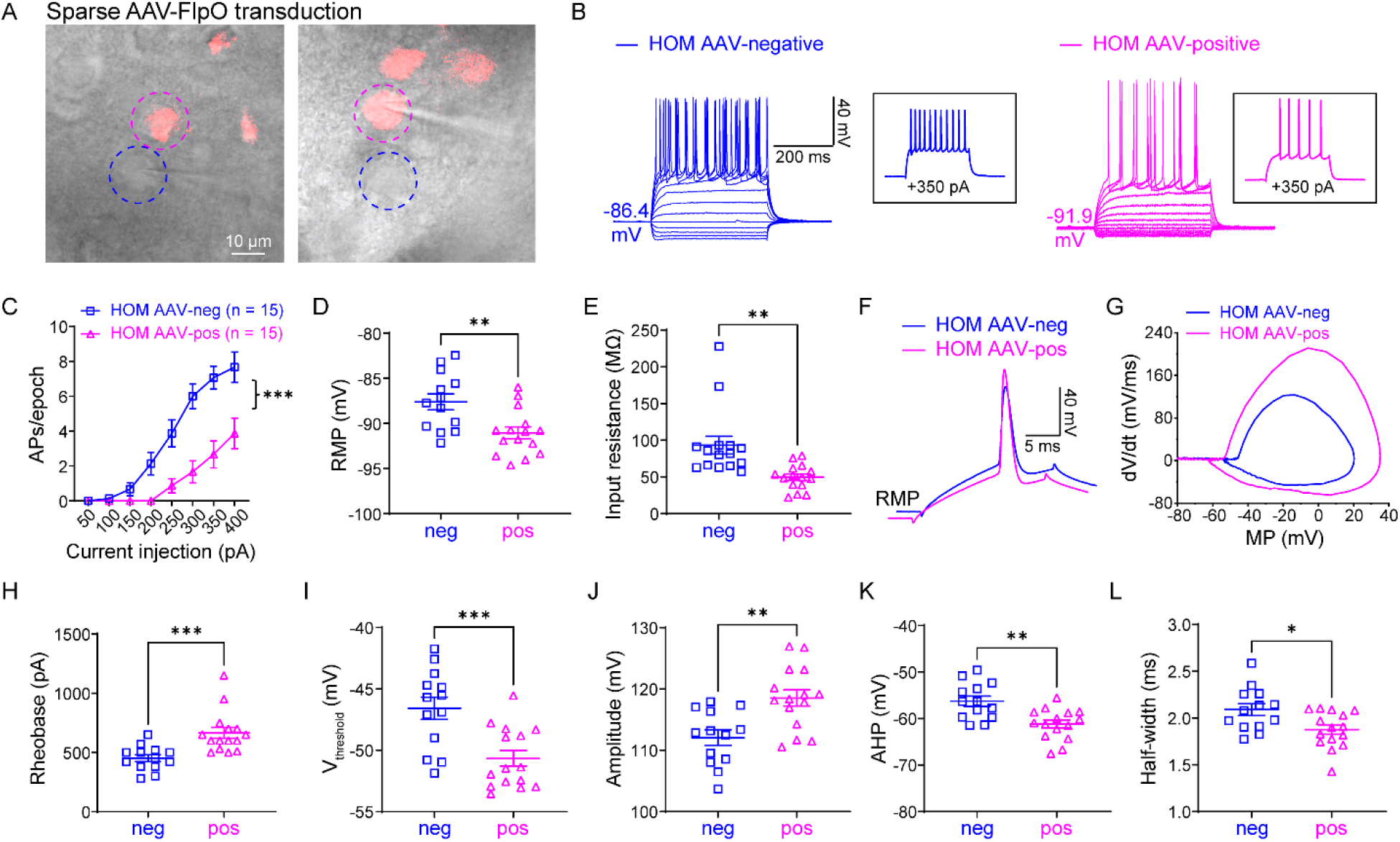
Elevated neuronal excitability is autonomous. (**A**) *Scn2a*^*gtKO/*gtKO^ (HOM) mice were injected with a dilute FlpO virus, transducing a subset of neurons in the striatum sparsely. Dashed circles highlight two neighboring AAV-negative (blue circle) and AAV-FlpO-positive (magenta circle) neurons. The images were taken in the cell-attached configuration, and after that, the target neurons were used for whole-cell recordings. (**B**) Representative current-clamp recordings of AAV-negative (blue) and AAV-FlpO-positive (magenta) MSNs in CPu of *Scn2a*^*gtKO/*gtKO^ mice were obtained at the RMP. A series of 400-ms hyperpolarizing and depolarizing steps in 50-pA increments were applied to produce the traces. Inset: representative trace in response to 350 pA positive current injection. (**C**) The average number of APs generated in response to depolarizing current pulses. Unpaired two-tailed non-parametric Mann-Whitney *U*-test for each current pulse: ***p < 0.001. (**D**) Individuals and average RMP values. Unpaired two-tailed Student’s *t*-test: **p < 0.01. (**E**) Individuals and average input resistance values at the RMP. Unpaired two-tailed Student’s *t*-test: **p < 0.01. (**F**) Typical spikes of MSNs with AAV-negative (blue) and with AAV-FlpO-positive (magenta) in HOM mice were obtained at the RMP. (**G**) Associated phase-plane plots. (**H-L**) Individuals and average spike rheobase, voltage threshold, amplitude, fAHP, and half-width values. Unpaired two-tailed Student’s *t*-test: *p < 0.05; **p < 0.01; ***p < 0.001. Data were shown as mean ± SEM.

### Downregulation of potassium channels contributes to the elevated action potential firings

To reveal the possible molecular basis underlying the enhanced neuronal excitability of *Scn2a*^*gtKO/gtKO*^ mice, we studied the gene expression profile using RNA sequencing (RNA-seq). We identified around nine hundred genes that were significantly up- or down-regulated in *Scn2a*^*gtKO/gtKO*^ mice compared to WT littermates (**Figure 4A**). *Scn2a* expression was at 29.6% of the WT value (**Figure 4B**), consistent with our qPCR (**Figure S4A**) and Western blot study (**Figure 2C and Figure S1C**). NaV1.6 and Na_v_1.2 are two major sodium channels often working in a coordinated fashion in principal neurons in the CNS, and the dysfunction of NaV1.6 is involved in seizures (*29–32*). In NaV1.6-deficient mouse models, Na_v_1.2 was upregulated, suggesting a compensatory relationship (*33, 34*). Interestingly, we detected a slightly reduced expression of NaV1.6 in *Scn2a*^*gtKO/gtKO*^ mice in our RNA-seq analysis. This reduction of NaV1.6 did not reach statistical significance (91.4±2.3% of WT, n = 4, p = 0.39) by qPCR validation (**Figure S4A**), indicating that our observed neuronal hyperexcitability is not likely to result from the compensation of the NaV1.6 channel expression.

**Fig. 4.**
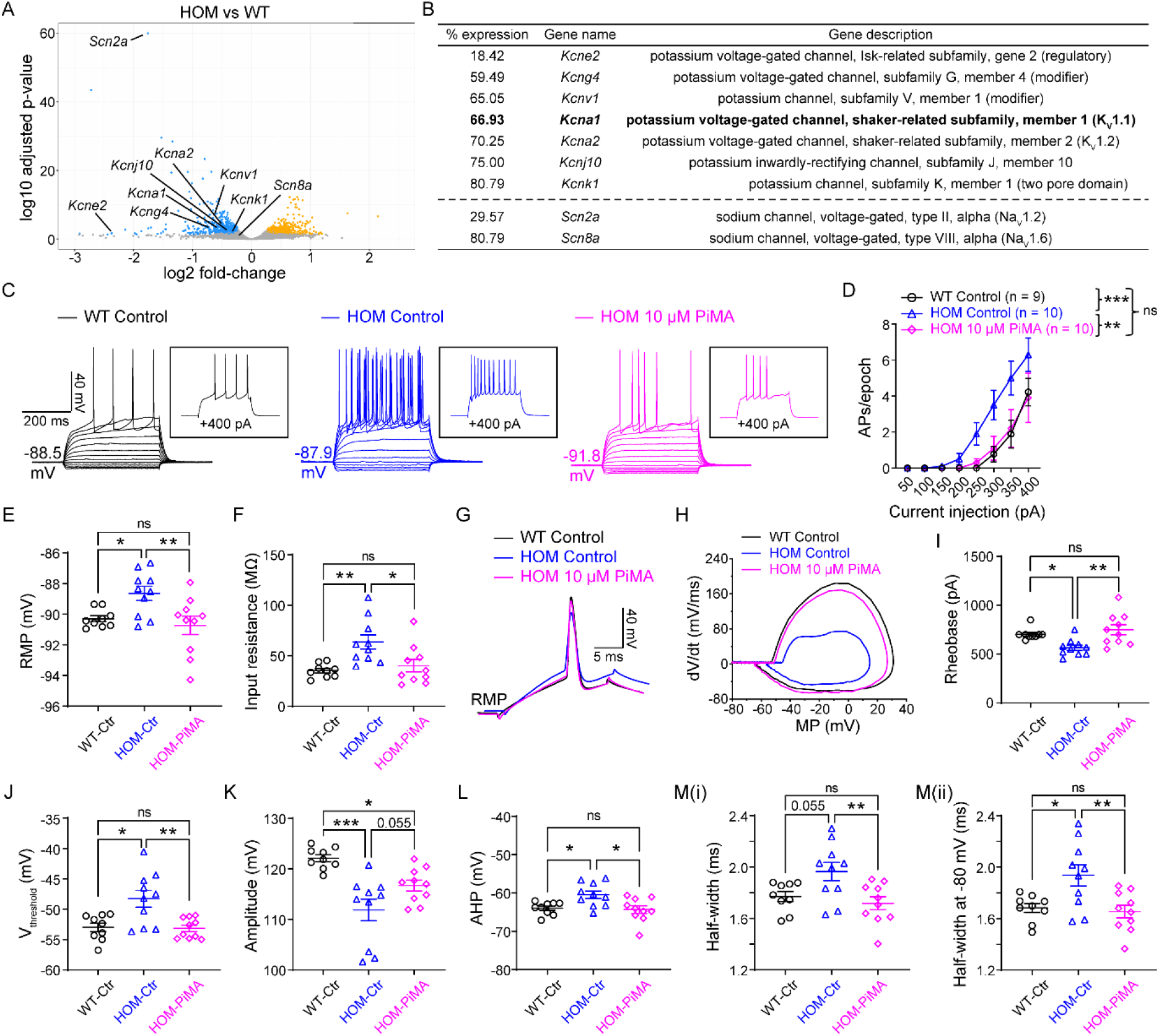
Activation of KV channels reverses elevated neuronal firings in adult Na_v_1.2-deficient mice. (**A**) Volcano plot displays *Scn2a* and *Scn8a*, as well all potassium channels that are statistically down-regulated in *Scn2a*^*gtKO/*gtKO^ (HOM) mice compared to WT mice identified by RNA-seq. Statistically significantly upregulated genes are shown in yellow and downregulated genes are shown in blue. (**B**) List of potassium channels that are significantly down-regulated in HOM mice compared to WT. (Hits that are identified from both DESeq2 and edgeR differential expression analysis with False Discovery Rate < 0.05 were listed). “% expression”: percentage expression of the gene in HOM mice considering the value of WT mice as 100%. (n = 4 mice for each group). (**C**) Representative current-clamp recordings of MSNs from WT slices perfused with 0.1% DMSO in aCSF (WT Control, black), HOM slices perfused with 0.1% DMSO in aCSF (HOM Control, blue), and HOM slices perfused with 0.1% DMSO in aCSF containing PiMA (HOM 10 μM PiMA, magenta) at the RMP. A series of 400-ms hyperpolarizing and depolarizing steps in 50-pA increments were applied to produce the traces. Inset: representative trace in response to 400 pA positive current injection. (**D**) The average number of APs generated in response to depolarizing current pulses at the RMP. Unpaired two-tailed non-parametric Mann-Whitney *U*-test for each current pulse: ns, no significance, *p > 0.05; **p < 0.01; ***p < 0.001. (**E**) Individuals and average RMP values. Unpaired two-tailed Student’s *t*-test: ns, no significance, *p > 0.05; *p < 0.05; **p < 0.01. (**F**) Individuals and average input resistance values at the RMP. Unpaired two-tailed Student’s *t*-test: ns, no significance, *p > 0.05; *p < 0.05; **p < 0.01. (**G**) Typical spikes of MSNs from WT slices perfused with 0.1% DMSO in aCSF (WT Control, black), HOM slices perfused with 0.1% DMSO in aCSF (HOM Control, blue), and HOM slices perfused with 0.1% DMSO in aCSF containing PiMA (HOM 10 μM PiMA, magenta) were obtained at the RMP. (**H**) Associated phase-plane plots. (**I-M**) Individuals and average spike rheobase, voltage threshold, amplitude, fAHP, and half-width values. Unpaired two-tailed Student’s *t*-test: ns, no significance, *p > 0.05; *p < 0.05; **p < 0.01; ***p < 0. 001. Data were shown as mean ± SEM.

Besides the NaV channels, potassium channels are also known to be major mediators setting the neuronal excitability (*35, 36*), and are often co-localized with NaV in the axon with high expression to regulate excitability (*37, 38*). Indeed, as the AP waveform was altered markedly in neurons with severe Na_v_1.2 deficiency, it is likely that the functions or expressions of potassium channels, which are responsible for many aspects of AP waveform, were disrupted in these neurons. Thus, we expanded our survey to include potassium channels. Notably, we found multiple potassium channel genes to be significantly downregulated (*Kcne2*, *Kcng4*, *Kcnv1*, *Kcna1*, *Kcna2*, *Kcnj10*, and *Kcnk1*) in our RNA-seq analysis (**Figure 4B**). As the top three genes are regulatory subunits or modifiers, we mainly focused on KV1.1 and KV1.2 (encoded by *Kcna1* and *Kcna2*, respectively), which are known to be involved in seizures (*39*). Our qPCR experiment validated that KV1.1 and KV1.2 were significantly downregulated (**Figure S4B**).

To understand the contribution of potassium channels towards neuronal excitability, we tested pimaric acid (PiMA, 10 μM) on MSNs in the brain slices of *Scn2a*^*gtKO/gtKO*^ mice. PiMA is a relatively general K channel opener but with demonstrated properties as a KV1.1-KV2.1 opener (*40*). While it might not be surprising that PiMA can affect neurons from WT mice, it was quite remarkable that PiMA almost completely rescued the excitability of MSNs from *Scn2a*^*gtKO/gtKO*^ mice to the WT range (**Figure 4C-M**). By pre-incubation of 10 μM PiMA for 10 min or more, we found that PiMA could significantly rescue the AP firings of MSNs from *Scn2a*^*gtKO/gtKO*^ mice towards the WT range (**Figure 4C-D**). Strikingly, most of the parameters, including input resistance, RMP, AP rheobase, voltage threshold, amplitude, fAHP, and half-width values were reversed towards the WT values in the presence of PiMA as well (**Figure 4E-M**).

As a selective KV1.1 opener (4-Trifluoromethyl-L-phenylglycine, 4TFMPG, 100 μM) was recently reported (*41*), we further investigated the role of the specific KV1.1 in MSNs of *Scn2a*^*gtKO/gtKO*^ mice. Notably, we found that the pre-incubation of 100 μM 4TFMPG for 10 min or more could significantly reverse the hyperexcitability of MSNs as well as the RMP and rheobase values of *Scn2a*^*gtKO/gtKO*^ mice (**Figure S4C-E, I, N-O, and T**). Interestingly, different from the relatively broad potassium channels opener (PiMA), 4TFMPG was not able to rescue the AP voltage threshold, amplitude, fAHP, half-width, or input-resistance (**Figure S4F-M** and **Figure S4Q-X**).

To further understand whether the change of expressions in these KV channels is due to Na_v_1.2 deficiency, we performed the qPCR experiment with striatal tissues from mice injected with AAV-FlpO, in which the expression of *Scn2a* was restored. Our data revealed that after the restoration of *Scn2a* expression by FlpO, the expressions of KV1.1 and KV1.2 were increased (**Figure S4B**), suggesting that neurons have a dynamic adaptation mechanism to regulate gene expression in responses to the change of *Scn2a* expression level. Taken together, our data indicate that KV channels (especially KV1.1) are important mediators of the hyperexcitability phenotypes observed in MSNs with severe Na_v_1.2 deficiency.

### *In vivo* neuronal firing in the CPu region is enhanced in adult Na_v_1.2-deficient mice

To test whether the enhanced neuronal excitability in brain slices translates into increased neuronal firing *in vivo*, we performed high-density *Neuropixels* recording on *Scn2a*^*gtKO/gtKO*^ and WT mice. The whole experiment pipeline consists of five steps, including surgery to implant headplate, recovery, recording, and postmortem imaging (**Figure S5A**). Mice were allowed to recover for 14 days before *in vivo* recording. The *Neuropixels* probe, consisting of ~300 recording electrodes across 3-mm length, was inserted into the CPu region to record the neuronal firing of mice in their resting-state (**Figure S5B-C**). After spike-sorting, units were manually verified and clear action potential waveform could be identified from both *Scn2a*^*gtKO/gtKO*^ and WT mice (**Figure S5D**). A published set of criteria were used to isolate putative MSNs (*42, 43*). Notably, our data demonstrated that putative MSNs from the CPu region of *Scn2a*^*gtKO/gtKO*^ mice display a higher mean firing frequency compared to WT mice (**Figure S5E**). Together, our data suggest that the neuronal hyperexcitability observed in brain slice recording can manifest as enhanced *in vivo* firings of head-fixed mice in their resting-state.

## DISCUSSION

Here in this paper, we report a counterintuitive finding that severe Na_v_1.2 deficiency renders hyperexcitability of principal MSNs in the striatum and pyramidal neurons in the mPFC, challenging the conventional paradigm. We further demonstrated that this hyperexcitability is reversible even in adult mice, showing a dynamic adaptive ability of neurons. Moreover, we provided evidence to suggest that the compensatory reduction in expressions of KV channels is a possible mechanism underlying this hyperexcitability, revealing a remarkable interplay between neuronal excitability and gene regulation. *In vivo* study further demonstrated that this elevated neuronal excitability identified in brain slices can be translated into enhanced neuronal firing in live mice. Our data thus provided a plausible explanation for the mysterious epileptic seizure phenotypes in humans with *SCN2A* deficiency, and identified molecular targets for potential therapeutic interventions.

Na_v_1.2 channel plays a variety of roles in the initiation, propagation, and backpropagation of APs during development and adulthood (*20, 44–47*). In the early stage of development, Na_v_1.2 is suggested to be the main sodium channel expressed in the axon initial segment (AIS) (*1, 48, 49*). Later in the development, NaV1.6 becomes the dominating channel in the axon and distal AIS, while the expression of Na_v_1.2 is re-distributed to other parts of the neurons including proximal AIS and dendrites (*20, 25, 47, 50*). A recent study found that pyramidal neurons of the mPFC from adult *Scn2a*^*+/−*^ mice have impaired excitatory postsynaptic current but intact AP firing (*20*). Here we revealed that severe Na_v_1.2 deficiency beyond a 50% reduction level in neurons surprisingly leads to hyperexcitability, which is an intrinsic property of the neurons that can be modulated in adulthood. However, how severe Na_v_1.2 deficiency changes neuronal excitability during early development remains to be determined. It is also worth noting that the expression of Na_v_1.2 in parvalbumin (PV) or somatostatin (SST) interneurons is limited (*5, 20*), and Na_v_1.2 does not seem to play a functional role in these interneurons (*20*). Nevertheless, it is still possible that while the expression of Na_v_1.2 in PV and SST interneurons is low, the severe reduction of *Scn2a* in principal neurons may result in compensatory adaptation, which indirectly affects the excitability of interneurons. It would be interesting to further explore these possibilities in a future study.

Because of the strong expression of *SCN2A* in principal neurons and its key roles to support AP firing, it is well accepted that increased Na_v_1.2 channel activity leads to enhanced excitability of principal neurons and aggravates seizures (*2, 12, 51*). Intriguingly, Na_v_1.2 deficiency, which is mainly found in ASD/ID cases, and conventionally expected to impair neuronal excitability, is also associated with epilepsies (*6, 10, 52*). It is estimated that 20~30% of ASD/ID patients with Na_v_1.2 deficiency develop “late-onset” seizures (*2, 11*). Treating epileptic seizures in these patients with Na_v_1.2 deficiency is extremely difficult, and the use of sodium channel blockers has been shown to exacerbate, rather than alleviate, the seizures (*11*). Our current data, together with published studies on *Scn2a*^*+/−*^ mice (*20*), may suggest a new paradigm. A moderate deficiency of Na_v_1.2 (i.e., loss-of-function variants) may impair neuronal excitability contributing to ASD and ID, whereas a severe deficiency of Na_v_1.2 (i.e., protein-truncating variants) tips the balance, resulting in neuronal hyperexcitability and increased seizure susceptibility. Notably, an independent study by the Bender lab (Spratt et al, co-submission) found that 100% knockout of *Scn2a* in a subset of pyramidal neurons in the mPFC results in hyperexcitability as well.

Potassium channels are known to play major roles in neuronal excitability and epileptic seizures (*53–57*). The AP waveform, which is highly influenced by the orchestration of a variety of potassium channels, is strongly disrupted in neurons with Na_v_1.2 deficiency, further suggesting the involvement of potassium channels. Our RNA-seq results identified multiple potassium channels to be significantly downregulated in *Scn2a*^*gtKO/gtKO*^ mice, including voltage-gated potassium channels (i.e., KV1.1 and KV1.2), as well as two-pore potassium channels (*Kcnk1*) and inward rectifier potassium channels (*Kcnj10*) (**Figure 4A-B**). KV1.1, for example, is abundantly expressed in principal neurons of the CNS, and contributes to the threshold as well as the interspike intervals during repetitive firing (*35*). Additionally, it is known that KV1.1 can form heteromultimeric channels with KV1.2 (*35*), which was identified to be downregulated in *Scn2a*^*gtKO/gtKO*^ mice by our RNA-seq analysis as well.

In this current study, we have obtained evidence to show that PiMA markedly reverses the elevated action potential firings associated with severe Na_v_1.2 deficiency (**Figure 4C-D**). PiMA is a relatively general potassium channels opener, with demonstrated properties as a KV1.1-KV2.1 and large-conductance Ca^2+^-activated K channel (BK) opener (*40*). It is worth noting that besides increased action potential firings, neurons from *Scn2a*^*gtKO/gtKO*^ mice display reduced action potential amplitude, higher voltage-threshold, increased input resistance, and elevated fAHP (**Figure 1** and **Figure S1**), which could be modulated by different potassium channels (*58, 59*). The reduced driving force on KV channels due to the change of AP waveform, including fAHP, might contribute to the hyperexcitability phenotypes as well. Nevertheless, PiMA turns out to rescue most of these altered parameters (**Figure 4**), which is unexpected but may indicate that PiMA has additional targets beyond KV1.1-KV2.1 and BK channels. Importantly, a relatively selective KV1.1 opener (4TFMPG) is also able to reduce neuronal hyperexcitability, suggesting that the enhanced action potential firing in *Scn2a*^*gtKO/gtKO*^ mice could be largely attributed to the KV1.1 channel (**Figure S4C-E, N-O**). Notably, 4TFMPG was not able to restore the input resistance, AP voltage threshold, amplitude, fAHP, or half-width values (**Figure S4F-M** and **Figure S4Q-X**), largely fitting the notion that 4TFMPG is selective for KV1.1 (*41*). Viral delivery of a specific gene is a different approach to elucidate the precise role of each distinct potassium channel towards neuronal excitability. Indeed, AAV-KV1.1 has been suggested as novel gene therapy to reduce seizures (*60, 61*). It would be appealing to test the effect of AAV-KV1.1 in *Scn2a*^*gtKO/gtKO*^ mice in a follow-up study. However, as several other potassium channels were also found to be downregulated, a multiple-gene delivery approach might be needed to deeply assess the contributions of this collection of ion channels toward neuronal hyperexcitability.

In summary, our results reveal an unexpected hyperexcitability phenotype in neurons with severe Na_v_1.2 deficiency, which is reversible and likely due to the compensatory reduction in expressions of potassium channels. The maladaptation or “over-compensatory” from potassium channels is likely a cause leading to the hyperexcitability of neurons to promote seizures. While it is a demonstrated clinical observation that patients with *SCN2A* deficiency often develop intractable seizures, there are no disease models that exist thus far for mechanistic investigation of this observation. Neuronal hyperexcitability identified in this unique Na_v_1.2-deficient mouse model is the first step towards the understanding of disease mechanisms underlying severe *SCN2A* deficiency. Our findings may explain the puzzling clinical observation that a portion of patients with Na_v_1.2 deficiency still develop seizures, and guide further development of interventions targeting KV channels to treat Na_v_1.2 deficiency-related disorders (*62*).

## MATERIALS AND METHODS

### Mouse strains

C57BL/6N-*Scn2a1*^*tm1aNarl*^/Narl (referred to as *Scn2a*^*WT/gtKO*^) mice were generated from the National Laboratory Animal Center Rodent Model Resource Center based on a modified gene-trap design (*23, 24*). The generation and basic characterization of this mouse model are available in our recent article (*22*). The targeting construct (tm1a trapping cassette) was electroporated into C57BL/6N embryonic stem cells, and founders in a pure C57BL/BN background were obtained to produce mice for experiments. All animal experiments were approved by the Institutional Animal Care and Use Committee (IACUC). Mice were same-sex housed in mixed-genotype groups (3-5 mice per cage) on vented cage racks with 1/8” Bed-o-cobb bedding (Anderson, Maumee, OH, USA) and > 8 g of nesting material as enrichment (shredded paper, crinkle-cut paper, and/or cotton nestles) on a 12hr light cycle. Food (2018S Teklad from Envigo) and reverse osmosis water were given *ad-lib*. Heterozygous (HET, *Scn2a*^*WT/gtKO*^) mice were used as breeding pairs to obtain homozygous (HOM, *Scn2a*^*gtKO/gtKO*^) mice and WT littermates for study. Whenever possible, investigators were blind to the genotype of the mice.

### Reagents

Reagents used were as follows: N-(2-aminoethyl) biotin amide hydrochloride (NEUROBIOTIN™ Tracer, SP-1120, from Vector Laboratories), Alexa 488-conjugated streptavidin (Molecular Probes, Eugene, OR, USA), Tetrodotoxin citrate (sodium channel blocker) was solubilized in pure water at a stock concentration of 500 μM); Pimaric acid (PiMA, KV channels opener) was solubilized in DMSO at a 1000× stock concentration of 10 mM; 4-(Trifluoromethyl)-L-phenylglycine (4TFMPG, KV1.1 specific opener) was solubilized in 1 M hydrochloric acid at a 1000× stock concentration of 100 mM; (2-Fluorophenyl) glycine (2FPG, KV1.1 specific opener) was solubilized in 0.25 M hydrochloric acid at a 1000× stock concentration of 100 mM.

### Antibodies

Primary antibodies used were: Rabbit anti-SCN2A (Na_v_1.2) (1: 1000, Alomone Labs, ASC-002), mouse anti-β-Actin (1:2000, Cell Signaling Technology, 3700S), and GAPDH (D16H11) XP^®^ Rabbit mAb (1:2000, Cell Signaling Technology, 5174S). Secondary antibodies were: IRDye® 680RD Goat anti-Rabbit IgG Secondary Antibody (1:5000, LI-COR Biosciences, AB_10956166) and IRDye® 680RD Goat anti-Mouse IgG Secondary Antibody (1:5000, LI-COR Biosciences, AB_10956588).

### Genotyping

Mice were labeled and genotyped via ear punch at weaning (21-28 days old). Genotyping for the tm1a cassette was performed using gene-specific polymerase chain reaction (PCR) on DNA extracted from ear tissues with a tissue DNA extraction kit (Macherey-Nagel, Bethlehem, PA, USA) with primers (forward 5’ to 3’: GAGGCAAAGAATCTGTACTGTGGGG, reverse: GACGCCTGTGAATAAAACCAAGGAA). The wild type allele’s PCR product is 240 base pairs (bp) and the tm1a (gtKO) allele’s PCR product is 340 bp.

### Adeno-associated virus (AAV) production

pAAV-EF1a-mCherry-IRES-Flpo was a gift from Karl Deisseroth (*63*) (Addgene plasmid # 55634 and viral prep # 55634-AAVrg; http://n2t.net/addgene:55634; RRID: Addgene_55634), AAV9-PHP.eB-EF1a-mCherry-IRES-Flpo with the titer of 2.56×10^13^ GC/mL was packed by Penn Vector Core (http://pennvectorcore.med.upenn.edu/); Control virus, PHP.eB-Ef1a-DO-mCherry-WPRE-pA with the titer of 1.2×10^13^ GC/mL was packed by Bio-Detail Corporation.

### Surgical procedures

For all surgeries (except as noted), mice were systemically anesthetized with ketamine and xylazine, and received analgesic buprenorphine to help postoperative recovery.

### AAV injections

For systemic delivery of virus, each adult mouse received 2×10^11^ infections of FlpO- or control AAV virus via tail vein injection. For viral injection into the brain to label neurons sparsely, mice were anesthetized with ketamine/xylazine (100/10 mg/kg, i.p.) and secured in a stereotaxic apparatus with ear-bars (RWD Ltd, China). After exposing the skull via a small incision, small holes for each hemisphere were drilled for injection based on coordinates to bregma. Mice were bilaterally injected with AAV virus (diluted into ~5×10^10^ infections units per mL with PBS) into the caudate nucleus and the putamen (CPu, dorsal striatum) (coordinates of the injection sites relative to bregma: AP +1.30 mm, ML ±1.25 mm, DV −3.30 mm; AP +0.50 mm, ML ±2.00 mm, DV −3.25 mm, 0.5-1 μL per point) and the nucleus accumbens (NAc, ventral striatum) (coordinates of the injection sites relative to bregma: AP +1.30 mm, ML ±1.25 mm, DV −4.50 mm, 0.5-1 μL per point) with sharpened glass pipettes (Sutter Instrument), self-made to have a bevel of 35° and an opening of 20-μm diameter at the tip (*64*), attached to syringe needles (200-μm diameter). The pipette was filled from the back end with mineral oil and attached to a syringe needle mounted in a microinjection syringe pump (World Precision Instruments, UMP3T-2). Before injection, the viral suspension was suctioned through the tip of the pipette. The skull over the target coordinates was thinned with a drill and punctured with the tip of the pipette. The pipette was inserted slowly (120 μm/min) to the desired depth. The virus was slowly (~100-150 nL/min) injected to the desired location. Before being retracted out of the brain, the pipette was left at the same place for 10 min when the injection was finished. The virus was allowed to express for at least three weeks before electrophysiological recordings. Animals were allowed to recover from surgery for one week and their body weight and health conditions were closely monitored during recovery. The accurate location of injection sites and viral infectivity were confirmed in all mice *post-hoc* by imaging of sections (50 μm in thickness) containing the relevant brain regions.

### Perfusions and tissue processing

For immunostaining, mice were administered an overdose of anesthesia and transcardiacally perfused with ice-cold PBS followed by 4% paraformaldehyde (PFA) (For *LacZ* staining, 4% PFA was replaced by 2% formaldehyde + 0.2% glutaraldehyde in PBS, hereinafter inclusive). After perfusion, brain slices were dissected out and post-fixed in 4% PFA overnight at 4°C. Tissues were cryoprotected by sinking in gradient sucrose (10%, 20%, and 30%) with 0.01 M PBS at 4°C and subsequently frozen in 20% sucrose and 30% sucrose in 1× phosphate-buffered saline (PBS) for 24-48 hrs. Samples were frozen in Optimal Cutting Temperature compound using dry ice and stored at −80°C. Tissue sections of 20 μm in thickness were taken on a cryostat (Leica CM1950) and allowed to air dry on slides, followed by analysis on a confocal microscope (Zeiss LSM 900 or Nikon A1R-MP).

### *LacZ* (β-galactosidase) staining

Both *Scn2a*^*gtKO/gtKO*^ and WT mice with or without AAV injection were processed at the same time under the same condition to minimize variation. Cryosections were fixed with 2% formaldehyde + 0.2% glutaraldehyde in PBS for 5 min. Then sections were washed at least 5 min in PBS (with 0.02% Triton X-100 for optimal reduction of unspecific binding of antibodies). Tissues were covered with a volume of freshly prepared staining solution [X-Gal solution added into Iron Buffer (1/19, v/v) and mixed thoroughly for 10 min], sufficient to fully cover the specimen (e.g., 50 μL) and incubate for 15-30 min at 37°C in a humid chamber until cells were stained blue. Color development was checked under a microscope and incubation time was continued if necessary. Specimen were washed three times with PBS and mounted in glycerol before storage after removing PBS. Images were analyzed under an upright light microscope.

### Immunostaining and imaging analysis

Cryosections (20 μm in thickness) were permeabilized, incubated in blocking buffer (0.5% Triton X-100 and 5% normal goat serum in PBS) for one hour at room temperature, and overlaid with primary antibodies overnight at 4°C. Then, the corresponding Alexa Fluor 488-, 594- or 647-conjugated secondary antibodies were applied. All stained sections were mounted with DAPI-containing mounting solution and sealed with glass coverslips. All immunofluorescence-labeled images were acquired using a confocal microscope (*65*).

### RNA sequencing

#### RNA extraction

Four *Scn2a*^*gtKO/gtKO*^ (HOM) and four WT littermate mice were used to extract RNA. Mice were given an overdose of anesthesia and transcardiacally perfused with ice-cold PBS. Acute coronal brain slices containing cortex and striatum (300-μm in thickness) were cut using a vibratome (Leica VT1200S, Germany). Cortex and striatum were rapidly microdissected, immersed into liquid nitrogen, and stored at −80°C until use (same procedures for Western Blotting and qPCR). Based on the manufacturer’s instructions, total RNAs were extracted with TRIzol reagent (Thermo Fisher Scientific, 15596018) from mouse cerebral tissues.

#### Library preparation and sequencing

Novogene prepared libraries using the TruSeq Stranded kit (Illumina, San Diego, CA) and RNA quality was assessed using an Agilent Nano RNA ChIP. Paired-end 150 bp reads were sequenced using the NovaSeq 6000.

#### Analysis

Reads were quality trimmed and Illumina TruSeq adapter sequences were removed using Trimmomatic v.0.36 (*66*). A sliding window approach to trimming was performed, using a window size of 5 and a required average Phred (quality) score of 16. Bases falling below a Phred score of 10 at the start and end of reads were trimmed and reads shorter than 20 bases in length after trimming were removed. FastQC v. 0.11.7 (*67*) was run to observe data quality before and after trimming/adapter removal. STAR v. 2.5.4b (*68*) was used to align reads to the Ensembl *Mus musculus* genome database version GRCm38.p6. The htseq-count script in HTSeq v.0.7.0 (*69*) was run to count the number of reads mapping to each gene. HTSeq used Biopython v.2.7.3 in the analysis. HTSeq was run utilizing the GTF file on “intersection-nonempty” mode. The HTSeq feature was set to “exon” and the attribute parameter was set to “gene_id” and the -- stranded=reverse option was set. The Bioconductor packages DESeq2 v.1.22.2 and edgeR 3.24.3 were used for differential expression analysis. Genes that were identified as differentially expressed in both packages were used as high confidence differentially expressed genes and were used in subsequent pathway analysis. The Benjamini-Hochberg false discovery rate correction was used to correct p-values for multiple testing. To improve power, low expression transcripts were filtered out of the data before performing differential expression analysis. The threshold chosen was to filter out all genes expressed at lower than 0.5 counts per million (CPM) in all samples combined. After filtering, 18,134 genes were remaining. The expression of genes between WT and HOM were deemed significant if the adjusted p-value < 0.05. The Bioconductor package biomaRt v. 2.38.0 was used to perform annotation of genes. ClusterProfiler v. 3.10.1 was used to perform pathway and gene ontology enrichment analysis.

### Western blotting

Brain tissues were homogenized in ice-cold RIPA lysis and extraction buffer (Thermo Fisher Scientific, 89901) supplemented with protease and phosphatase inhibitors (Thermo Fisher Scientific, A32953), sonicated, and cleared by centrifugation (10,000× g, 10 min, at 4°C). Protein concentration in the supernatant was determined by (determined by Nanodrop, Thermo Scientific). Proteins in 1× sample buffer [62.5 mM Tris-HCl (pH 6.8), 2% (w/v) SDS, 5% glycerol, 0.05% (w/v) bromophenol blue] were denatured by boiling at 95°C for 5 min. For each sample, 40 μg total proteins were loaded to the 8% sodium dodecyl sulfate-polyacrylamide (SDS-PAGE) gels and transferred onto PVDF membrane (Millipore, IPFL00010) by electrophoresis. Blots were blocked in 5% nonfat milk in Tris-buffered saline and Tween 20 (TBST) for 1 h at room temperature and probed with the primary antibody in 5% milk-TSBT overnight at 4°C. After overnight incubation, the blots were washed three times in TBST for 15 min, followed by incubation with corresponding IRDye® 680RD secondary antibodies in TBST for 2h at room temperature. Following three cycles of 15 min washes with TBST, the immunoreactive bands were scanned and captured by the Odyssey® CLx Imaging System (LI-COR Biosciences) and quantitatively analyzed by densitometry with Image Studio Lite 5.2 (LI-COR Biosciences) or ImageJ software (NIH). Each sample was normalized to its β-actin or GAPDH, then normalized with the corresponding WT littermates.

### RNA isolation, reverse transcription, and qPCR analysis

Total RNAs were extracted with TRIzol reagent (Thermo Fisher Scientific, 15596018) from mouse cerebral tissues according to the manufacturer’s instructions. 4 μg RNA was subjected to reverse transcription (RT) with a Maxima First Strand cDNA Synthesis Kit (Thermo Fisher Scientific, K1672). The resulting cDNAs were subjected to quantitative PCR analysis using the PowerUp™ SYBR™ Green Master Mix (Thermo Fisher Scientific, A25777) and specific primers in a C1000 Touch PCR thermal cycler (Bio-Rad). *Gapdh* and *β-actin* mRNA levels were used as an endogenous control for normalization using the ΔCt method (*70*). In brief, test (T):ΔCt^T^ = [Ct^T^ (target gene) - Ct^T^ (internal control)]; Amount of the target = 2^−ΔCt^.

### Patch-clamp recordings

#### Acute slice preparations

Electrophysiology was performed in slices prepared from 2-5 months-old *Scn2a*^*gtKO/gtKO*^ and corresponding control mice. Mice were deeply anesthetized with ketamine/xylazine (100/10 mg/kg, i.p., 0.1 mL per 10 grams of body weight), and then transcardially perfused, and decapitated to dissect brains into ice-cold slicing solution containing the following (in mM): 110 choline chloride, 2.5 KCl, 1.25 NaH_2_PO_4_, 25 NaHCO_3_, 0.5 CaCl_2_, 7 MgCl_2_, 25 glucose, 0.6 sodium ascorbate, 3.1 sodium pyruvate (bubbled with 95% O_2_ and 5% CO_2_, pH 7.4, 305-315 mOsm). Acute coronal slices containing PFC and/or striatum (300-μm in thickness) were cut by using a vibratome (Leica VT1200S, Germany) and transferred to normal artificial cerebrospinal fluid (aCSF) (in mM): 125 NaCl, 2.5 KCl, 2.0 CaCl_2_, 2.0 MgCl_2_, 25 NaHCO_3_, 1.25 NaH_2_PO_4_, 10 glucose (bubbled with 95% O_2_ and 5% CO_2_, pH 7.4, 305-315 mOsm). Then, slices were incubated at 37°C for 20-30 minutes and stored at room temperature before use. Slices were visualized under IR-DIC (infrared-differential interference contrast) using a BX-51WI microscope (Olympus) with an IR-2000 camera (Dage-MTI).

#### *Ex vivo* electrophysiological whole-cell recordings

All somatic whole-cell patch-clamp recordings were performed from identified striatal MSNs or mPFC layer V pyramidal neurons. The selection criteria for MSNs were based on morphological characteristics with medium-sized cell body presenting polygon or diamond viewed with a microscope equipped with IR-DIC optics (BX-51WI, Olympus), and numerous dendritic spines and their hyperpolarized RMP (lower than −80 mV) based on published method (*71*). Layer V pyramidal cells with a prominent apical dendrite were visually identified mainly by location, shape, and pClampex online membrane test parameters. Putative pyramidal cells in layer 5b were identified based on regular spiking characteristics (*20, 72, 73*). To minimize variability, recordings were made on cells with low or high HCN expression levels, corresponding to intratelencephalic (IT) or pyramidal tract (PT) neurons, respectively. The selection criterion for PT pyramidal cells was based on their firing properties and shape of the AP (i.e., all cells’ intrinsic ability to generate, upon subthreshold depolarization possessed a prominent after-hyperpolarization and significant membrane-potential sags induced by both hyperpolarizing and depolarizing current injection at the soma). Recordings of PT neurons were used for further analysis.

For whole-cell current-clamp recordings, the internal solution contained (in mM): 122 KMeSO_4_, 4 KCl, 2 MgCl_2_, 0.2 EGTA, 10 HEPES, 4 Na_2_ATP, 0.3 Tris-GTP, 14 Tris-phosphocreatine, adjusted to pH 7.25 with KOH, 295-305 mOsm. The sag ratio, input resistance, and firing number were obtained in response to a series of 400 ms current steps from −200 pA to +400 pA in increments of 50 pA, each sweep duration of 5 s with cells held at the normal RMP or a fixed potential of −80 mV. The sag ratio was calculated with the equation:

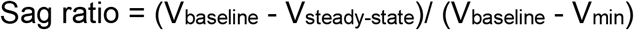

Where V_baseline_ is the resting membrane potential or −80 mV, V_min_ is the minimum voltage reached soon after the hyperpolarizing current pulse, and V_steady-state_ (V_ss_) is the voltage recorded at 0-10 ms before the end of the −200 pA stimulus.

The input resistance was calculated with the equation:

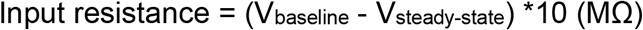

Where V_baseline_ is the resting membrane potential or −80 mV, and V_steady-state_ (V_ss_) is the voltage recorded at 0-10 ms before the end of the −100 pA stimulus.

The RMP, AP threshold, amplitude, fast afterhyperpolarization (fAHP), and half-width values were obtained in response to a 20 ms current step of the smallest current to obtain an intact AP, each sweep duration of 1.5 s and start-to-start intervals of 10 s with cells held at the normal RMP or a fixed potential of −80 mV. The RMP, AP threshold, amplitude, fAHP, and half-width values were analyzed using the Clampfit 11.1 inbuilt statistics measurements program (Criteria as the baseline, peak amplitude, antipeak amplitude, and half-width). The threshold was defined as the Vm when dV/dt measurements first exceeded 15 V/s.

We used thin-wall borosilicate pipettes (BF150-110-10) with open-tip resistances of 3-5 MΩ. All recordings were started at least 1 min after breakin to stabilize the contact between the glass electrode and the cell membrane, and finished within 10 min to avoid large voltage changes due to the internal solution exchange equilibrium. Recordings were performed with an Axon MultiClamp 700B amplifier (Molecular Devices) and data were acquired using pClamp 11.1 software at the normal RMP or a fixed potential of −80 mV, filtered at 2 kHz and sampling rate at 20 kHz with an Axon Digidata 1550B plus HumSilencer digitizer (Molecular Devices). Slices were maintained under continuous perfusion of aCSF at 32-33°C with a 2-3 mL/min flow. In the whole-cell configuration series resistance (Rs) 15-30 MΩ, and recordings with unstable Rs or a change of Rs > 20% were aborted.

To study the effect of KV channels openers, the 1000× stocks were freshly diluted with aCSF, respectively. After 10 min perfusion of each opener or the corresponding vehicle control (0.1% DMSO in aCSF for PiMA, and aCSF for 4TFMPG), the target neurons were studied with the continuous perfusion of the chemicals. One or two neurons were patched for each brain slice, and recordings were discarded if a slice was perfused with KV channels openers for more than 30 min.

For cell labeling, the internal solution contains 0.1-0.2% (w/v) neurobiotin tracer. At the end of the electrophysiological recording (about 30 min), slices were treated as previously described (*74*). Briefly, sections were fixed in 4% paraformaldehyde in 0.1 M phosphate buffer (pH 7.4) for 20-30 min at room temperature, and subsequently washed 3-4 times for 30 min in 0.1 M phosphate-buffered saline (PBS, pH 7.4) at 4°C. Sections were then incubated in Alexa 488-conjugated streptavidin (overnight at 4°C, 1: 250 in blocking solution) to visualize neurobiotin.

### *Neuropixels* recordings and data analysis

#### Surgeries

Animal preparation was performed as described previously (*64, 75*). Mice were anesthetized and head-fixed and underwent stereotaxic surgery to implant a metal headframe with a 10-mm circular opening (Narishige, MAG-1, and CP2) for head-fixation. An incision was made over the skin. The skin and periosteum were removed, and a thin layer of cyanoacrylate (Krazy glue) was applied to attach the headplate and cover the exposed skull. A layer of clear Stoelting™ Dental Cement (Fisher Scientific, 10-000-786) was then applied on top of cyanoacrylate and forms a chamber around the skull to contain the ground wire and aCSF during electrophysiological recordings. The animals received two weeks recovery period after surgery before commencing experiments. Before electrophysiological recordings, a 600 μm diameter craniotomy was prepared to access the intended brain regions with *Neuropixels* probes.

#### *In vivo* recordings

Electrophysiological recordings were made with *Neuropixels* probes in head-fixed mice. On the day of the experiment, the mouse was placed under light isoflurane anesthesia. A ground wire was secured to the skull, and the exposed brain was covered with a layer of 4% agar in aCSF. Following recovery from anesthesia, the mouse was head-fixed on the experimental rig. Before insertion, the probe tip was painted with CM-DiI. Briefly, the *Neuropixels* probe was secured to an arm of stereotaxic, and the backside of the probe was dipped into a 1 μL droplet of CM-DiI (Thermo Fisher Scientific, C7000) dissolved in ethanol (1 μg/μL). The ethanol was allowed to evaporate, and the CM-DiI was dried onto the backside of the tip. The probe was then inserted slowly (120-480 μm/min) into the striatum (coordinates of the injection sites relative to bregma: AP +1.30 mm, ML ±1.25 mm, 4.50 mm in depth) through the craniotomy on the skull. After reaching the desired depth for a probe, the probe was allowed to settle for 10 minutes before the commencement of recording. The first 384 electrodes were turned on in the *Neuropixel* probe, which corresponds to about 3.8 mm length probe. At the end of each recording session, the probe was retracted out of the brain and cleaned using Tergazyme (Alconox) followed by washing with distilled water. The probe insertion was verified by identifying the DiI fluorescence in sectioned brain tissue.

#### Data acquisition and Analysis

All data were acquired with a 30-kHz sampling rate under the Open Ephys GUI (https://open-ephys.atlassian.net/wiki/spaces/OEW/pages/963280903/Neuropix-PXI). A 300-Hz high-pass filter was present in the Neuropixels probe, and another 300-Hz high-pass filter (3rd-order Butterworth) was applied offline before spike sorting.

Spike waveforms were automatically extracted from the raw data using Kilosort 2.0 (https://github.com/MouseLand/Kilosort/releases/tag/v2.0). The outputs were loaded into PHY (*76*) for manual refinement, which consisted of merging and splitting clusters, as well as marking non-neural clusters as “noise”. Noise units were identified by their abnormal waveform shape, as well as distinct cyclical patterns in the autocorrelogram. A set of heuristic rules based on the features of waveforms to remove abnormal waveforms [the parameters were used for this purpose were peak-to-trough (PT) ratio < 0.99 and recovery slope < 0]. Waveforms for each unit were extracted from the raw data, and then averaged. All the averaged waveforms were used to calculate the mean waveform.

Striatal single units were classified according to the methods described previously (*42*), using mean firing rate, mean waveform peak width at half-maximum, mean waveform trough width at half-minimum, and ISI distribution. These values were averaged across epochs when a cell was present in multiple epochs. The standard classification for the clusters was defined as follows: fast-spiking interneurons (FSIs): firing rate > 3 Hz, peak width < 0.2 ms, and a ratio of trough width to peak width (TPR) < 2.7 (TPR was estimated by k-means clustering and was more reliable than exact trough width for FSIs); tonically-active neurons (TANs): < 5% of ISIs less than 10 ms, a median ISI > 100 ms, and peak width (0.2-0.35 ms) and trough width (0.1-0.2 ms) above the 95^th^ percentile for the remainder of the units; unclassified units had low TPR and/or narrow trough widths (< 0.3 ms) but firing rates < 2 Hz; all other units were considered putative medium spiny neurons (MSNs).

### Quantification and statistical analysis

Normality and variance similarity was measured by GraphPad Prism before we applied any parametric tests. Two-tailed Student’s *t*-test (parametric) or unpaired two-tailed Mann-Whitney U-test (non-parametric) was used for single comparisons between two groups. Other data were analyzed using one-way or two-way ANOVA with Tukey correction (parametric) or Kruskal-Wallis with Dunn’s multi comparison correction (non-parametric) depending on the appropriate design. *Post hoc* comparisons were carried out only when the primary measure showed statistical significance. Error bars in all figures represent mean ± SEM. p values less than 0.05 were considered statistically significant. Statistical significance of differences at p < 0.05 is indicated as one asterisk (*), p < 0.01 is indicated as two asterisks (**), and p < 0.001 is indicated as three asterisks (***) in all figures. Mice with different litters, body weights, and sexes were randomized and assigned to different treatment groups, and no other specific randomization was used for the animal studies.

## SUPPLEMENTARY MATERIALS

Supplementary material for this article is available online at TBD.

## General

We thank Dr. Amy Brewster and Dr. Chongli Yuan for the critical reading of this paper.

## Funding

This work is supported by Purdue startup funding, Ralph W. and Grace M. Showalter Research Trust, and Purdue Big Idea Challenge 2.0 on Autism to Y.Y. The authors gratefully acknowledge support from the Purdue University Institute for Drug Discovery and Institute for Integrative Neuroscience. M.E. is supported by the National Science Foundation (NSF) Graduate Research Fellowship Program (GRFP) (DGE-1842166). Yang lab is grateful to the *FamilieSCN2A* foundation for the Action Potential Grant support. This project was funded, in part, with support from the Indiana Clinical and Translational Sciences Institute funded, in part by Award Number UL1TR002529 from the National Institutes of Health, National Center for Advancing Translational Sciences, Clinical and Translational Sciences Award. The Yang lab appreciates the bioinformatics support of the Collaborative Core for Cancer Bioinformatics (C^3^B) with support from the Indiana University Simon Comprehensive Cancer Center (Grant P30CA082709), Purdue University Center for Cancer Research (Grant P30CA023168), and Walther Cancer Foundation. The content is solely the responsibility of the authors and does not necessarily represent the official views of the National Institutes of Health.

## Author contributions

J.Z., X.C., Y.Y. designed the experiments. J.Z., X.C., M.E., S.L., A.P., T.S.A., J.W., Z.M., Z.Q., J.L., T.X., Y.L., Y.W., M.I.O.A., N.A.L. performed the experiments and analyzed the data. J.A.S, K.J., Z.H., N.A.L., W.C.S. participate in data analysis and experimental design. Y.Y. supervised the project. J.Z. and Y.Y. wrote the paper with inputs from all authors.

## Competing interests

The authors declare no competing interests.

## Data and materials availability

Additional results can be found in the supplemental material. The data that supports the findings of this study are available from the corresponding author upon reasonable request.

---

**Figure S1.**
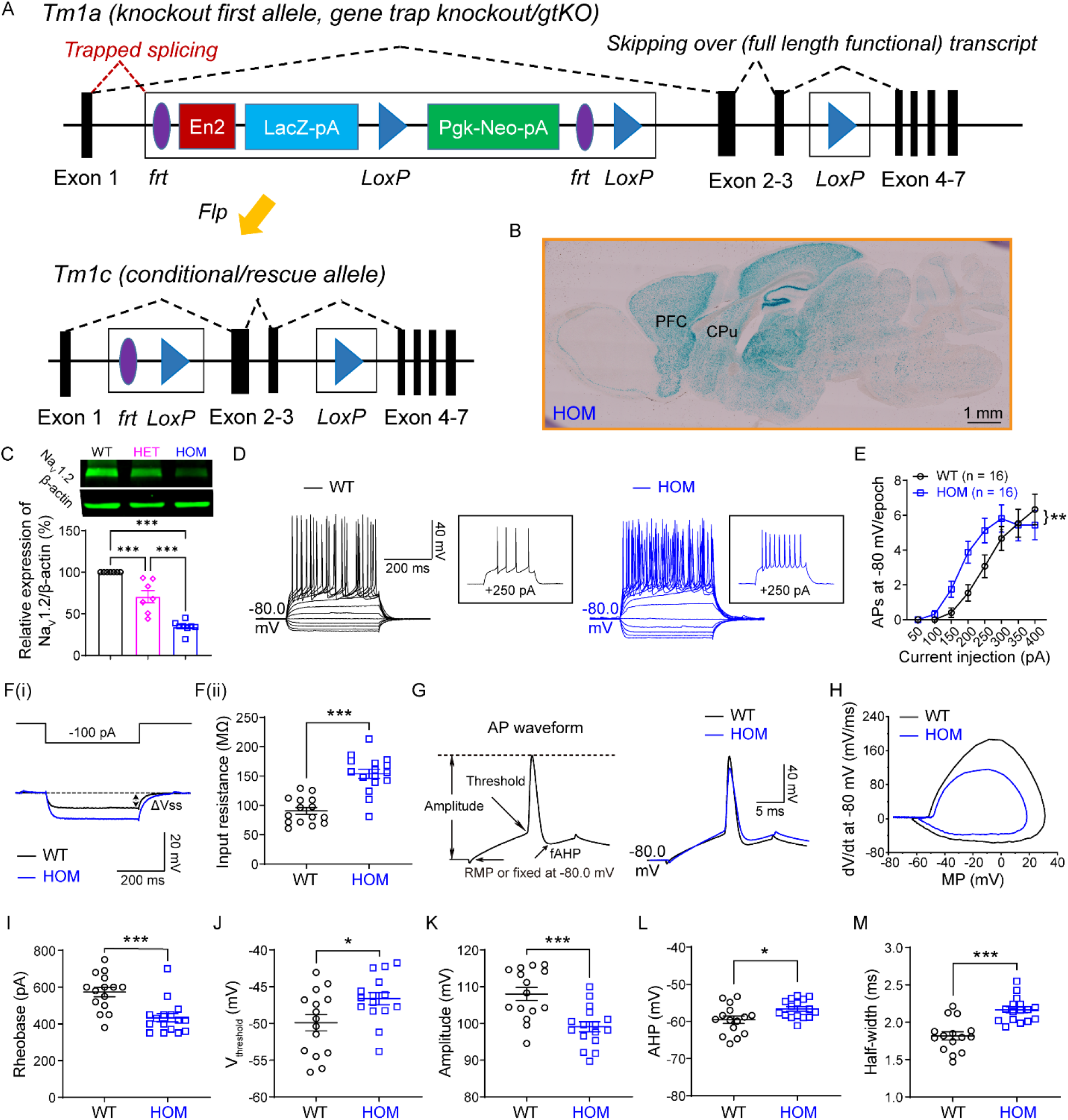
Elevated neuronal firings of striatal MSNs at a fixed membrane potential of −80 mV in adult Na_v_1.2-deficient mice. Related to Figure 1. (**A**) gtKO allele has an inserted tm1a trapping cassette between the Exon 1 and Exon 2 of *Scn2a* gene in the genome, which traps the transcription from Exon 1 to tm1a cassette, resulting in “gene-trap” knockout of *Scn2a*. In the presence of Flp recombinase, frt sites flanked trapping cassette will be removed, producing conditional (“rescue”) allele that allows the expression of *Scn2a* at the WT level. *frt*, *Flp* recognition target (purple); *En2*, engrailed-2 splice acceptor (red); *LacZ*, *lacZ* β-galactosidase (light blue); *LoxP*, locus of X-over P1 (dark blue); and *Neo*, neomycin (green). (**B**) gtKO cassette contains a *LacZ* element and is driven by the native *Scn2a* promoter. Thus, the *LacZ* expression can be used as a surrogate of *Scn2a* expression. Representative *LacZ* staining of a sagittal slice from a *Scn2a*^*gtKO/gtKO*^ (HOM) mouse showing a strong blue signal across the brain including the prefrontal cortex (PFC) and dorsal striatum (CPu, caudate nucleus and the putamen). (**C**) Upper: Representative Western blots of striatal tissues from WT (black circle), HET (magenta diamond), and HOM (blue square) mice. Lower: associated quantification of Na_v_1.2 protein. One-way ANOVA followed by Tukey’s multiple-comparison test: ***p < 0.001. (**D**) Representative current-clamp recordings of MSNs from WT (black) and HOM (blue) mice were obtained at a fixed membrane potential of −80 mV. A series of 400-ms hyperpolarizing and depolarizing steps in 50-pA increments were applied to produce the traces. Inset: representative trace in response to 250 pA positive current injection. (**E**) The average number of APs generated in response to depolarizing current pulses at −80 mV. Unpaired two-tailed non-parametric Mann-Whitney *U*-test for each current pulse: **p < 0.01. (**Fi**) Representative traces in response to 100 pA negative current injection. V_steady-state_ (V_ss_) is the voltage recorded at 0-10 ms before the end of the stimulus. (**Fii**) Individuals and average input resistance values at −80 mV. Unpaired two-tailed Student’s *t*-test: ***p < 0.001. (**G**) Left: plot of a typical AP showed its various phases. Right: typical spikes of MSNs from WT (black) and HOM (blue) mice were obtained at a fixed membrane potential of −80 mV. (**H**) Associated phase-plane plots. (**I-M**) Individuals and average spike rheobase, voltage threshold, amplitude, fAHP (fast after-hyperpolarization), and half-width values. unpaired two-tailed Student’s *t*-test: *p < 0.05; ***p < 0.001. Data were shown as mean ± SEM.

**Figure S2.**
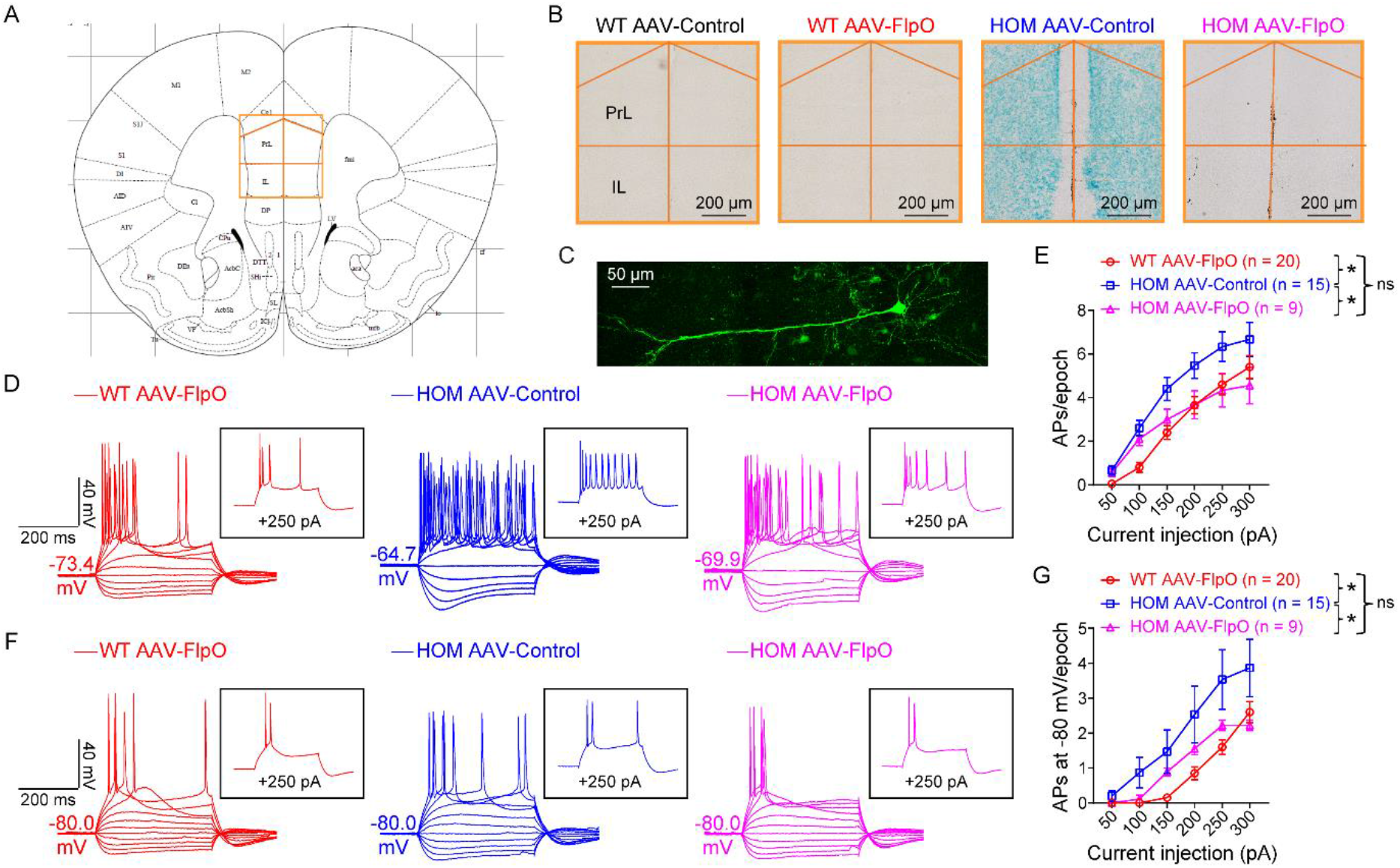
Elevated neuronal firings of layer V pyramidal cells in the mPFC are reversible by FlpO-mediated rescue in adult Na_v_1.2-deficient mice. Related to Figure 2. (**A-B**) *LacZ* staining of coronal brain slices containing mPFC from WT and *Scn2a*^*gtKO/gtKO*^ (HOM) mice, which were systemically administered with AAV-Control or AAV-FlpO. PrL, prelimbic cortex; IL, infralimbic cortex. (**C**) A typical layer V pyramidal neuron in the mPFC was labeled by neurobiotin. Scale bar, 50 μm. (**D**) Representative current-clamp recordings of pyramidal cells from WT mice transduced with AAV-FlpO (red), HOM mice transduced with AAV-Control (blue), and HOM mice transduced with AAV-Control (magenta) at the RMP. A series of 400-ms hyperpolarizing and depolarizing steps in 50-pA increments were applied to produce the traces. Inset: representative trace in response to 250 pA positive current injection. (**E**) The average number of APs generated in response to depolarizing current pulses at the RMP. Unpaired two-tailed non-parametric Mann-Whitney *U*-test for each current pulse: ns, no significance, p > 0.05; *p < 0.05. (**F**) Representative current-clamp recordings of layer V pyramidal cells in the mPFC from WT transduced with AAV-FlpO (red), HOM transduced with AAV-Control (blue) and HOM transduced with AAV-Control (magenta) at a fixed membrane potential of −80 mV. A series of 400-ms hyperpolarizing and depolarizing steps in 50-pA increments were applied to produce the traces. Inset: representative trace in response to 250 pA positive current injection. (**G**) The average number of APs generated in response to depolarizing current pulses at −80 mV. Unpaired two-tailed non-parametric Mann-Whitney *U*-test for each current pulse: ns, no significance, p > 0.05; *p < 0.05. Data were shown as mean ± SEM.

**Figure S3.**
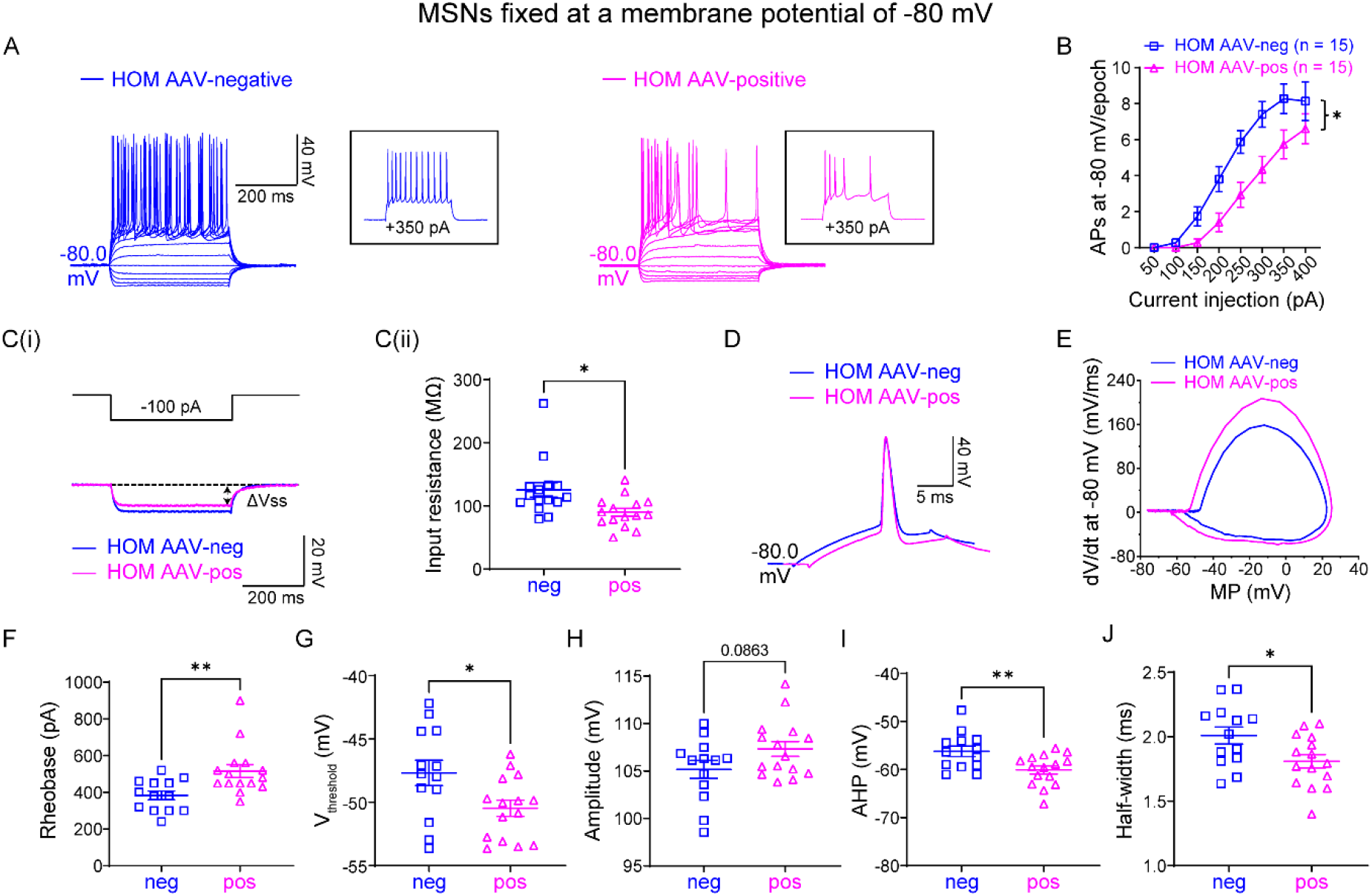
*Ex vivo* recordings of MSNs at a fixed membrane potential of −80 mV in adult Na_v_1.2-deficient mice with a dilute AAV-FlpO-mCherry injection. Related to Figure 3. (**A**) Representative current-clamp recordings of MSNs with AAV-negative (blue) and with AAV-FlpO-positive (magenta) in *Scn2a*^*gtKO/gtKO*^ (HOM) mice were obtained at a fixed membrane potential of −80 mV. A series of 400-ms hyperpolarizing and depolarizing steps in 50-pA increments were applied to produce the traces. Inset: representative trace in response to 350 pA positive current injection. (**B**) The average number of APs generated in response to depolarizing current pulses. Unpaired two-tailed non-parametric Mann-Whitney *U*-test for each current pulse: *p < 0.05. (**Ci**) Representative traces in response to 100 pA negative current injection. V_steady-state_ (V_ss_) is the voltage recorded at 0-10 ms before the end of the stimulus. (**Cii**) Individuals and average input resistance values at −80 mV. Unpaired two-tailed Student’s *t*-test: *p < 0.05. (**D**) Typical spikes of MSNs with AAV-negative (blue) or AAV-FlpO-positive (magenta) in HOM mice were obtained at a fixed membrane potential of −80 mV. (**E**) Associated phase-plane plots at −80 mV. (**F-J**) Individuals and average spike rheobase, voltage threshold, amplitude, fAHP, and half-width values. Unpaired two-tailed Student’s *t*-test: ns, no significance, p > 0.05; *p < 0.05; **p < 0.01. Data were shown as mean ± SEM.

**Figure S4.**
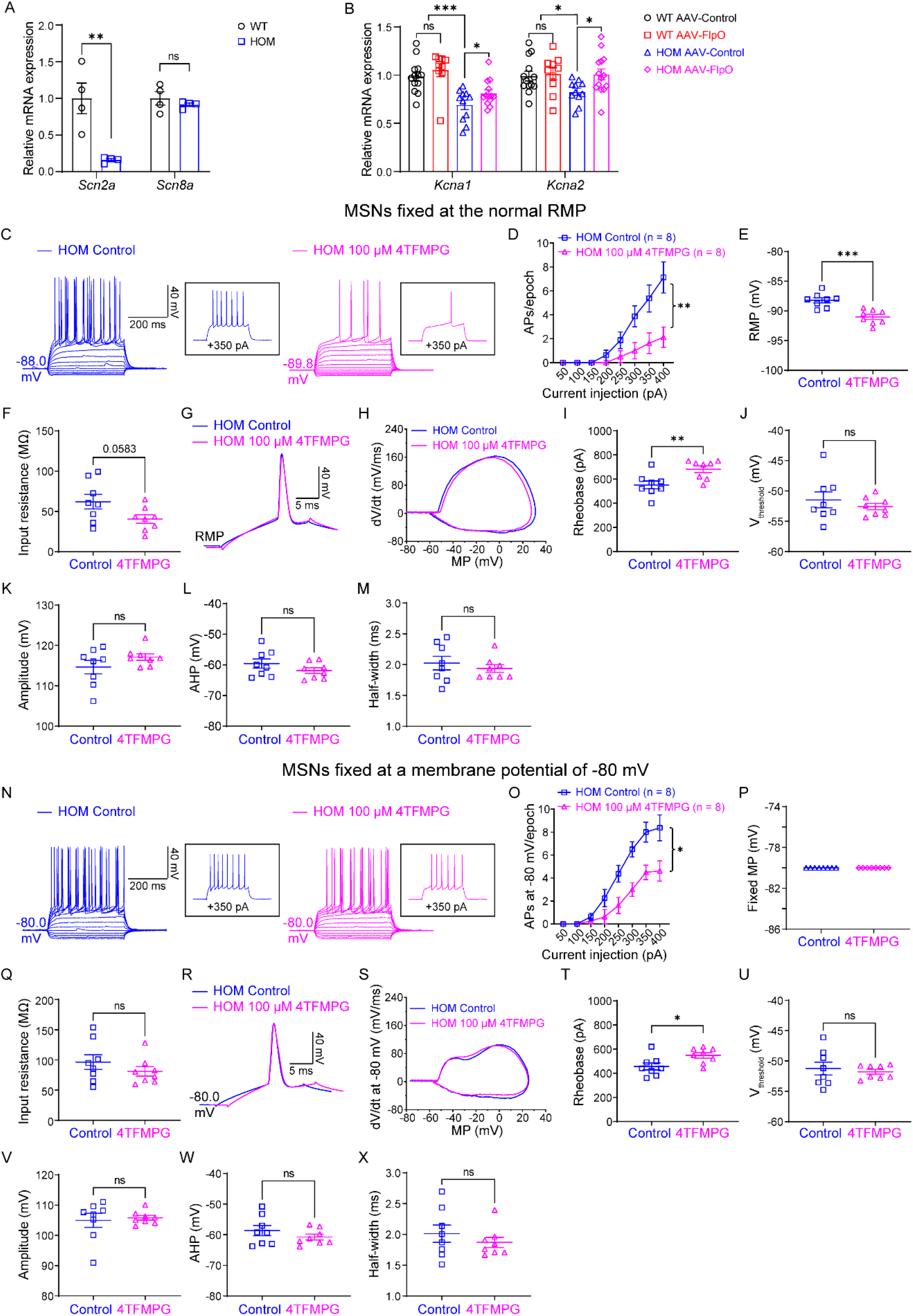
Specific activation of KV1.1 channel by 4TFMPG reverses the elevated neuronal firings in adult Na_v_1.2-deficient mice. Related to Figure 4. (**A**) Quantitative (q)PCR analysis of *Scn2a* and *Scn8a* mRNA in the striatum samples from WT and *Scn2a*^*gtKO/*gtKO^ mice. Unpaired two-tailed Student’s *t*-test for each group: ns, no significance, *p > 0.05; **p < 0.01. (**B**) qPCR analysis of *Kcna1* and *Kcna2* mRNA in the striatum samples from WT and HOM mice transduced with AAV-Control or AAV-FlpO, showing that the downregulated mRNA levels of KV1.1 and KV1.2 were reversible by FlpO-mediated restoration of Na_v_1.2 expression in adult Na_v_1.2-deficient mice. Unpaired two-tailed Student’s *t*-test: ns, no significance, *p > 0.05; *p < 0.05; ***p < 0.001. (**C**) Representative current-clamp recordings of MSNs from HOM slices perfused with aCSF (HOM Control, blue) and HOM slices perfused with aCSF containing 4TFMPG (HOM 100 μM 4TFMPG, magenta) at the RMP. A series of 400-ms hyperpolarizing and depolarizing steps in 50-pA increments were applied to produce the traces. Inset: representative trace in response to 350 pA positive current injection. (**D**) The average number of APs generated in response to depolarizing current pulses at the RMP. Unpaired two-tailed non-parametric Mann-Whitney *U*-test for each current pulse: **p < 0.01. (**E**) Individuals and average spike RMP values. Unpaired two-tailed Student’s *t*-test: ***p < 0.001. (**F**) Individuals and average input resistance values at the RMP. Unpaired two-tailed Student’s *t*-test: p = 0.0583. (**G**) Typical spikes of MSNs from HOM slices perfused with aCSF (HOM Control, blue) and HOM slices perfused with aCSF containing 4TFMPG (HOM 100 μM 4TFMPG, magenta) were obtained at the RMP. (**H**) Associated phase-plane plots. (**I-M**) Individuals and average spike rheobase, voltage threshold, amplitude, fAHP, and half-width values. (**N**) Representative current-clamp recordings of MSNs from HOM slices perfused with aCSF (HOM Control, blue) and HOM slices perfused with aCSF containing 4TFMPG (HOM 100 μM 4TFMPG, magenta) at a fixed membrane potential of −80 mV. A series of 400-ms hyperpolarizing and depolarizing steps in 50-pA increments were applied to produce the traces. Inset: representative trace in response to 350 pA positive current injection. (**O**) The average number of APs generated in response to depolarizing current pulses at −80 mV. Unpaired two-tailed non-parametric Mann-Whitney *U*-test for each current pulse: *p < 0.05. (**P**) fixed MP values for recording. (**Q**) Individuals and average input resistance values at −80 mV. Unpaired two-tailed Student’s *t*-test: ns, no significance, *p > 0.05. (**R**) Typical spikes of MSNs from *Scn2a*^*gtKO/gtKO*^ slices perfused with aCSF (HOM Control, blue) and *Scn2a*^*gtKO/gtKO*^ slices perfused with aCSF containing 4TFMPG (HOM 100 μM 4TFMPG, magenta) were obtained at a fixed membrane potential of −80 mV. (**S**) Associated phase-plane plots. (**T-X**) Individuals and average spike rheobase, voltage threshold, amplitude, fAHP, and half-width values. Unpaired two-tailed Student’s *t*-test: ns, no significance, *p > 0.05; *p < 0.05. Data were shown as mean ± SEM.

**Figure S5.**
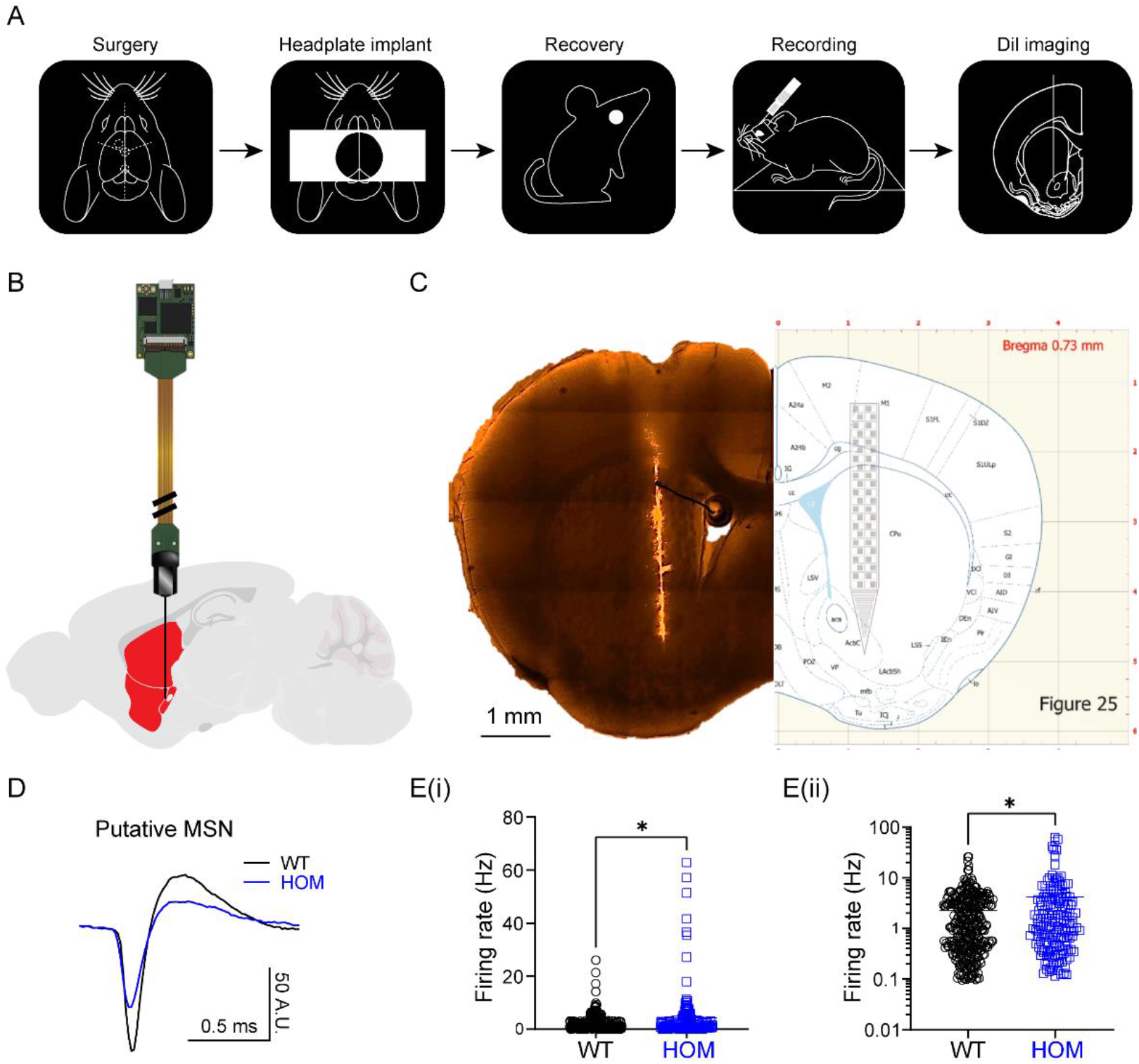
Elevated *in vivo* neuronal firings of putative striatal MSNs in adult Na_v_1.2-deficient mice. (**A**) Cartoon icons showing 5 steps in the *Neuropixels* recording experiment pipeline. (**B**) A cartoon illustration of a *Neuropixels* probe inserted into the striatum. (**C**) DiI staining of *Neuropixels* probe after recording in the mouse brain matched with brain map (the right panel was adapted from Figure 25 in the *Paxinos and Franklin’s The Mouse Brain in Stereotaxic Coordinates*). (**D**) Representative spike waveforms of *Neuropixels* recordings from putative MSNs of WT (black) and HOM (blue) mice. (**Ei**): Firing rate of putative MSNs of WT and HOM mice. (**Eii**): y-axis in log scale to show the firing rate of putative MSNs. n = 3 mice for each genotype; unpaired two-tailed Welch’s *t*-test: *p < 0.05. Data were shown as mean ± SEM.

## Notes

### Competing Interest Statement

The authors have declared no competing interest.

